# *Pseudomonas aeruginosa* shows between and within strains heterogeneity in virulence phenotype after passive exposure to zebrafish (*Danio rerio*) and nematodes (*Caenorhabditis elegans*)

**DOI:** 10.1101/2023.11.27.568077

**Authors:** Daniella Azulai, Anielka Espinoa, Maracela Talamantes, Heather L. Bennett, Gabriel G. Perron

**Affiliations:** Department of Biology, Reem-Kayden Center for Science and Computation, Bard College, Annandale-On-Hudson, NY 12504; Department of Biology, Trinity College, Hartford, CT, 06106; Center for Environmental Sciences and Humanities, Bard College, Annandale-On-Hudson, NY, 12504; Center for Systems Biology and Genomics, New York University, New York, NY, 10003

**Keywords:** Host-pathogen interactions, virulence, zebrafish, *C. elegans*, nematode, model system, opportunistic infection, *Pseudomonas aeruginosa*

## Abstract

The rapid emergence of diseases and parasites in aquatic wildlife requires improved methodologies to identify and characterize new and future pathogens. While microinjection of pathogens directly into a sentinel organism such as zebrafish enables the exploration of infection and immune response in the host, such methodology focuses primarily on identifying causative agents in events of aquatic wildlife mortality due to acute infection. Here, we present an updated protocol of infection by static immersion in larval zebrafish to investigate the possible effect of prolonged environmental exposure to an opportunistic pathogen. By controlling microbial growth and monitoring mortality over five days, we show that static immersion can detect minute differences in virulence profiles between and within different strains of *Pseudomonas aeruginosa*, an important opportunistic pathogen of animals and humans. We then conducted two sets of passive exposure virulence assays in *Caenorhabditis elegans*, an alternative model. We demonstrated the virulence phenotype, while showing slight differences between experimental models, showed similar trends. We believe that passive exposure thus offers a practical host-pathogen model that simulates opportunistic infection occurring in the environment and enables the detection of minute changes in virulence between and within bacterial strains.

## 1. Introduction

The rapid emergence of diseases and parasites affecting wildlife organisms is of increasing concern ^1^. While parasites can be a natural part of healthy ecosystems ^2^, environmental disturbance and climate change increased parasitic load in many environments ^3,4^. Such outbreaks of infectious diseases are especially problematic in organisms for which environmental changes can directly impact physiological responses, such as fishes ^5,6^ and insects ^7,8^. In addition to hindering conservation efforts ^9^, the increased burden of parasites in wildlife increases the risk of introducing novel pathogens in human populations ^10^.

The emergence of novel pathogens in any host depends on multiple factors. For example, environmental stressors such as antimicrobials can impact host health and immunity ^11,12^, creating a novel ecological virulent niche for bacteria that would otherwise not cause infection. Another possible factor is the development of genotypic and phenotypic diversity in a population of non-pathogenic bacteria that can lead to the evolution of virulence traits necessary to infect a new host ^13^. Such phenomena, following population source-sink dynamics ^14^, are more common when bacteria reach higher population densities in the environment, increasing the mutation supply rate ^15,16^. In this context, it is crucial to study the genotypic and phenotypic diversity in environmental reservoirs to predict the emergence of novel pathogens ^17,18^. Yet, the study of natural populations of bacteria remains sparse ^5,19^.

*Pseudomonas aeruginosa* is a striking example of an opportunistic pathogenic bacterium that can successfully invade a range of hosts and new hosts ^20^. The aquatic and soil bacterium can infect various organisms such as plants, nematodes, fruit flies, wax moths, zebrafish, and mammals under certain conditions ^21^. The bacterium is also associated with high rates of morbidity and mortality in intensive care unit (ICU) patients. Indeed, even within a single host like humans, *P. aeruginosa* can invade various tissues and sites ^22–24^. *P. aeruginosa’s* pathogenicity is due to various virulence factors and antibiotic resistance determinants, providing the bacterium with metabolic flexibility and great adaptability ^25,26^.

To understand just how the phenotypic heterogeneity found in *P. aeruginosa* contributes to virulence in animals and humans, it is crucial to identify reliable model systems that support a variability of phenotypic traits. Zebrafish, or *Danio rerio*, a well-established model organism whose immune system presents striking similarities to the immune systems of humans and other vertebrates ^27,28^, is an ideal candidate for identifying causative agents in aquatic wildlife diseases ^5^. While microinjection of pathogens is the preferred methodology in this host ^29^, passive exposure via static immersion is more characteristic of a large number of fish pathogens, including *P. aeruginosa*, transmitted via contact with the eyes, gills, skin, or even the gut following ingestion ^30,31^. Interestingly, previous work showed that *P. aeruginosa* strain PAO1, a common lab strain, did not yield mortality in larval zebrafish following a short exposure (*i.e.,* ∼ 5 hours) but triggered genes associated with a stress response ^32^.

Another model important model for the study of study host-pathogen interactions is *Caenorhabditis elegans*, a small nematode with a shorter developmental frame that is also susceptible to human pathogens ^33^. Indeed, the soil nematode used as a model system enabled the identification of various virulence-related genes in *P. aeruginosa* via transcriptional changes in *C. elegans* exposed to the bacterium ^34–36^. Interestingly, *C. elegans* predominantly feeds on bacteria cells, thus providing a different mode of entry for pathogenic bacteria.

Here, we present an investigation of *P. aeruginosa’s* heterogeneity in virulence phenotypes when exposed to two model systems, zebrafish embryos and *C. elegans*. By controlling microbial growth and monitoring mortality over a prolonged time against different strains of the bacterium, we show that passive exposure not only confirms that *P. aeruginosa* can cause mortality in zebrafish and soil nematodes but that the methodology can reveal important differences in virulence ability between the different strains of the bacterium.

## 2. Materials and Methods

### 2.1 Pseudomonas aeruginosa *strains*

We selected five *P. aeruginosa* strains from a more extensive collection at Bard College (Table 1). Clinical *P. aeruginosa* strains used in this study were chosen to reflect the diversity of phenotype observed in the bacterium, including different infection sites, country, and date of isolation. More information for each strain is available elsewhere ^21,24^.

**Table 1.** List of *Pseudomonas aeruginosa* strains.

| Strain Name | Country | Date Isolated | Source |
| --- | --- | --- | --- |
| A10 | France | 1882-1918 | Wound |
| AGO4092 | Greece | 1994 | Urine |
| ESP06B | Belgium | 1993 | Clinical |
| Lw1047 | Congo | 2001 | Blood |
| PA01 | Australia | 1955 | Wound |

To control for potential differences in virulence phenotype due to changes in growth between the strains of *P. aeruginosa*, we started each passive exposure assay with the same concentration of bacteria. While *P. aeruginosa* strains were grown on LB plates in *C. elegans* assays, the bacterium was maintained in 1x E3 media (i.e., 5mM NaCl, 0.17 mM KCl, 0.33 mM CaCl2, 0.33 mM MgSO4; pH 7.2) in zebrafish static immersion assays. For the latter, we measured bacterial growth by inoculating the bacterium at three concentrations (*e.g.,* 10^3^CFU/mL, 10^5^ CFU/mL, and 10^9^ CFU/mL) in E3 medium. The bacteria were incubated at 28 °C, and growth was monitored at 12 and 24 hours using serial dilution. Finally, to test whether *P. aeruginosa* subsisted in E3 media due to leftover LB nutrients following overnight culture, we compared the growth of *P. aeruginosa* starved in M9 media for 8 hours to that of *P. aeruginosa* washed in sterile 1x E3 media ^37^. In addition, both treatments were grown in either 1x E3 media or 1x E3 media containing 1% LB to test the effect of added nutrients on bacterial growth directly.

### 2.2 Zebrafish husbandry and static immersion virulence assay

Wild-type Tu zebrafish were maintained at the Bard College Biology Department’s zebrafish facility. The fish were kept on a 14/10-hr light/dark cycle in standard recirculating rack water maintained at 28°C with pH ranging from 7.0-7.4. Fish mating was performed in standard mating reservoirs. Upon release, embryos were cleaned by immersion in a 0.5% bleach solution for five minutes, followed by deionized water for five minutes, then the former steps were repeated. After washing, embryos were transferred to 100 mm^2^ Petri dishes in densities of 30-50 embryos and suspended in sterile 1x E3 media. 75% of the media was changed daily while embryos remained in the petri dish.

Larval zebrafish were then used to conduct a static immersion virulence assay adapted from van Soest et al. ^32^. While the latter exposed the larvae to a high concentration (10^9^ CFU/mL) of *P. aeruginosa* for five hours, we exposed the animals to a lower concentration (10^5^ CFU/mL) of *P. aeruginosa* for 72 hours. *P. aeruginosa* populations were prepared by centrifuging an overnight culture grown in LB at 37 °C, then pelleted before being resuspended in filter-sterilized 1x E3 media at the desired concentration. We placed one 3-day-post-fertilization (3-pdf) larvae in one well of a 24-well plate, with each well containing 2 mL of the bacterial mixture. All assays were conducted in a well-oxygenated closed environment at 28 °C for 72 hours. Fish mortality was then monitored every 12 hours by observing larval movement and confirmed by touching immobile larvae with a sterile pipette tip. As a control, we also monitored the mortality of 24 larvae growing in 2 mL of sterile 1x E3 media, and each assay was conducted twice with independent broods. All methods described herein concur with humane animal testing standards and were approved by the Bard Institutional Animal Care and Use Committee (IACUC; approval ID “Perron 2018”).

### 2.4 Maintenance of Caenorhabditis elegans and passive exposure virulence assay

Wild type (WT), or N2, worms were obtained from the *Caenorhabditis* Genetics Center. Animals were cultured and maintained with standard methods ^38^. *C. elegans* were grown on nematode growth medium (NGM) plates seeded with the *Escherichia coli* strain, OP50, at 20 °C. To synchronize worms by age, animals were bleached using a standard protocol (Porta-de-la-Riva et al. 2012), where gravid adults are dissolved in a buffer solution containing 4% sodium hypochlorite and 5M sodium hydroxide. The *C. elegans* embryos that survived were then washed with M9 buffer twice and allowed to hatch on NGM plates again seeded with OP50 at 20 °C. Two days later, L4 worms were washed three times with M9 buffer before being used in subsequent assays.

For our passive exposure virulence assay, we used the slow-killing (SK) plates described elsewhere ^33,35^. Approximately 30 triple-washed and synchronized L4 worms were plated onto clean SK plates seeded with the standardized liquid culture of bacteria. Worms were kept at 25 °C, and every two days, living worms were transferred onto freshly seeded SK plates such that they were repeatedly exposed to the same bacterial strain. We monitored mortality by prodding immobile *C. elegans* every 24 hours for seven days following the first exposure to the bacteria. If any *C. elegans* were unaccounted for, they were recorded as missing unless found within the next day.

### 2.6 Microscopy

We used microscopy to track the possible presence of *Paramecium* in the E3 medium, a common contaminant in the zebrafish experimental environment. Microscope images were obtained using a Nikon Eclipse E600, at 4x magnification. Fish were placed onto a glass microscope slide suspended within a drop of E3 media using a sterile pipette. Fluorescence images were obtained using the Nikon Eclipse E600 at 4x magnification equipped with the UV-1A filter cube attachment. While we sometimes observed a *Paramecium* bloom 24 hours following the death of larvae, we did not detect any important *Paramecium* populations during the survival stage of the experiment.

### 2.7 Statistical analyses

To investigate the effect of growth conditions of *P. aeruginosa* growing in 1x E3 media, we used an analysis of variance to compare growth after 24 hours measured as CFU/mL between populations initially inoculated with 10^5^ and 10^8^ CFU/mL. We then compared bacterial growth after 24 hours of bacteria population inoculated with 10^5^ CFU previously grown in LB or starved in M9 salt for 8 hours before inoculation.

To test for heterogeneity in bacterial virulence, we compared animal survival for both zebrafish and *C. elegans* using the Kaplan-Meier estimate of survival. The latter was implemented in the “survival” package of R ^39^, where each death is weighted using a log-rank test, and missing animals are given a value of 0 when they are not found. Finally, to test for a possible “batch” effect, we compared the two independent trials conducted for each model system using the Mann-Whitney test, a non-parametric test comparing the sum rank of two groups, and we compared the two independent assays performed for each bacterial strain-model system pair. All statistical analyses and model assumptions were tested in R 3.5.2 ^40^. Graphs were plotted using ggplot2 ^41^.

## 3. Results

### 3.1 Virulence in zebrafish

In order to establish a standard protocol to study bacteria virulence in immersion assay, we first investigated the growth conditions of *Pseudomonas aeruginosa* in the test medium. To do so, we monitored the growth of replicate populations of the bacterium inoculated at two different concentrations (*i.e.,* 10^5^ and 10^8^) over 24 hours. While the inoculation size had a significant effect on population growth (*F*_(1,4)_ = 31.85; *P* = 0.005), we observed that all the populations increased in size, reaching 6.42 • 10^8^ (4.51 • 10^7^) CFU/mL and 1.67 • 10^9^(1.55 10^8^) CFU/mL, respectively. Crucially, we also established that the bacterium’s growth was not due to residual nutrients stored in the bacterial cells or the inoculum due to prior growth in a nutrient rich medium (*F*_(3,8)_ = 0.0092; *P* = 0.99). These results suggest that *P. aeruginosa* can sustain a stable population in E3 media, providing an opportunity for virulence development. Based on our observation, we established that optimal static immersion conditions would be inoculation with 10^5^, thus allowing for stable bacterial populations in the medium.

We then tested whether long-term passive exposure to bacteria could result in mortality in larval zebrafish. To do so, we investigated the survival of zebrafish larvae exposed to five *P. aeruginosa* strains chosen to reflect the phenotypic diversity in the taxa. For each strain, we exposed 24 independent larval zebrafish randomly selected from a unique brood to identical bacterial growth conditions via static immersion in 2 mL microcosms. We then tracked larval mortality every 12 hours. Overall, we found that all five *P. aeruginosa* strains induced increased larvae mortality compared to a single mortality event in the control fish population (χ^2^_5_ = 99; *P* < 0.0001; Figure 1A-E).

**Figure 1.**
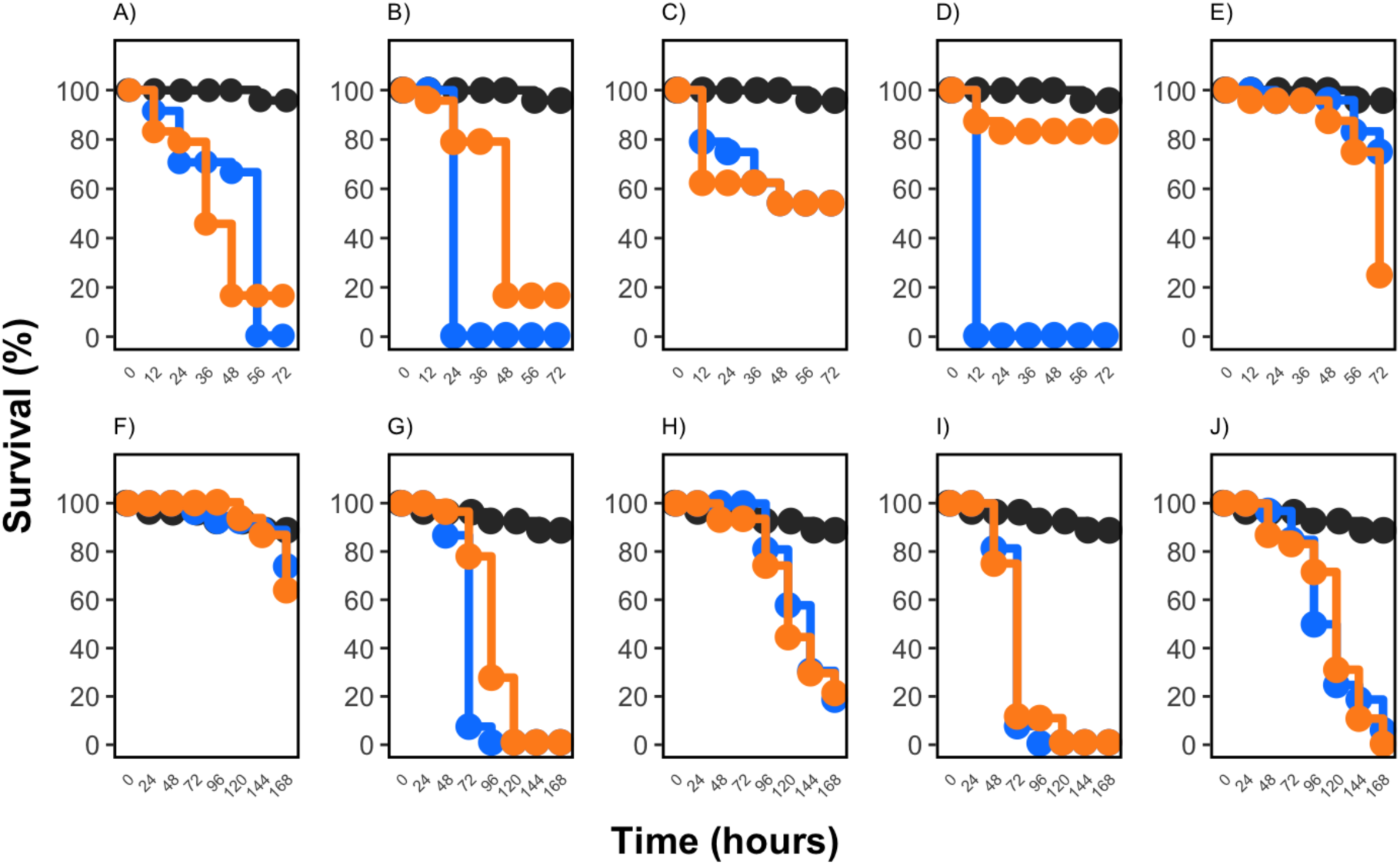
Survival curves of larval zebrafish and L4 *C. elegans* exposed to different *P. aeruginosa* strains. For zebrafish larvae, survival was computed every 12 hours from populations of 24 zebrafish larvae exposed to *P. aeruginosa* strain A) A10; B) AGO4092; C) ESP06B; D) Lw1047; and E) PA01, shown in blue (trial 1) and in orange (trial 2) and compared to a control population of 24 fishes not exposed to the bacterium (show in gray). For *C. elegans*, survival of 30 *C. elegans* exposed to *P. aeruginosa* strain F) A10; G) AGO4092; H) ESP06B; I) Lw1047, and J) PA01, shown in blue (trial 1) and orange (trial 2) and is compared to L4 worms exposed to OP50 (gray). For all tests, differences in survival were compared to control populations, and between each trial was tested using the Kaplan-Meier log-rank test.

Interestingly, we found that the *P. aeruginosa* strains induced mortality at varying rates. More specifically, we found that zebrafish exposed to strains A10 (χ^2^ _1_ = 49.3; adj-*P* < 0.001; Figure 1A) and AGO4092 (χ^2^ _1_ = 41.2; adj-*P* < 0.001; Figure 1B) displayed high mortality relatively early in the assay, with more than 80% larval zebrafish dying by 48 hours post-exposure. Larvae exposed to strain PAO1 (χ^2^ _1_ = 19.5; adj-*P* < 0.001; Figure 1E) also suffered high mortality, but at a slower rate than the aforementioned strains, with more than 80% larvae dying at 72 hours post-exposure, correlating with the development of mouth opening in the zebrafish larvae at about five days post fertilization. Fish exposed to strain ESP06B also showed lower mortality rates (χ^2^ _1_ = 13.1; adj-*P* = 0.0015; Figure 1C), but still significantly more than the control. Finally, fish exposed to strain Lw1047 did not show significant mortality compared to controls (χ^2^ _1_ = 0.9; adj-*P* > 0.99; Figure 1D).

While strains A10 and AGO4092 were not statistically different from each other (χ^2^ _1_ = 6.5; adj-*P* = 0.1), they differed from all other strains (adj-*P*s < 0.05). In addition to being statistically different from strains A10 and AGO4092, strain PAO1 was statistically different from strain Lw1047 (χ^2^ _1_ = 11.3; adj-*P* < 0.001) but not strain ESP06B (χ^2^ _1_ = 0.2; adj-*P* > 0.99). These results suggest that we observed three phenotype classes among the *P. aeruginosa* strains in zebrafish larvae: high (80% mortality), low (50% mortality), and absent.

To investigate the reliability of our results, we repeated the above static immersion assay with a second independent larvae brood. Again, we exposed populations of 24 independent larval zebrafish randomly selected from a unique brood to identical bacterial growth conditions via static immersion in 2 mL microcosms for each strain. While we observed the same overall trends with strains A10 and AGO4092 showing high mortality and ESP06B showing the lowest larval mortality, survival curves were often significantly different for each strain between the two trials. When comparing the two trials, we found a statistical difference in survival in fishes exposed to strain A10 (χ^2^ _1_ = 23.2; adj-*P* < 0.001; Figure 1A); AGO4092 (χ^2^ _1_ = 50.4; adj-*P* < 0.001; Figure 1B); Lw1047 (χ^2^ _1_ = 92.6; adj-*P* < 0.001; Figure 1D); and PAO1 ( χ^2^ _1_ = 11.8; adj-*P* < 0.001; Figure 1E). This second trial confirmed the consistent presence of three virulence phenotypes but also demonstrated an important source of variability between trials.

### 3.2 Virulence in nematodes

We then investigated whether *P. aeruginosa* would show a similar virulence profile in *C. elegans*. To do this, we fed independent populations of synchronized L4 nematodes on either of five different bacterial strains as described above. Control populations were fed on OP50, the standard laboratory bacterial diet. Survival for each nematode was scored every 24 hours. Overall, we found that every *P aeruginosa* strain, except for A10 (χ^2^ _1_ = 0.4; adj-*P* > 0.99; Figure 1F), increased mortality compared to *C. elegans* fed the control diet. Moreover, we found that *C. elegans* survival was significantly lower when exposed to strain AGO4092 (χ^2^ _1_ = 47; adj-*P* = 0.002; Figure G), ESP06B (χ^2^ _1_ = 23; adj-*P* < 0.001; Figure H), Lw1046 (χ^2^ _1_ = 30.4; adj-*P* < 0.001; Figure I), PAO1 (χ^2^ _1_ = 30.3; adj-*P* < 0.001; Figure J). We also found that strains AGO4092 and Lw1047, while significantly different from each other (χ^2^ _1_ = 29.4; adj-*P* < 0.001; Figure G) and had the highest mortality rate early in the assay, with only 6.67% and 0% of the exposed *C. elegans* surviving the first 72 hours. Strains ESP06B and PA01, which did not significantly (χ^2^ _1_ = 4.8; adj-*P* = 0.15), still led to 90% and 96% mortality, respectively, by the end of our experiment.

Again, to investigate the reliability of our results, we repeated the above assays with a second independent trial. Unlike our assays with the zebrafish model systems, we found that most bacterial strains tested showed similar virulence patterns over the two trials. Indeed, strain A10 (χ^2^ _1_ = 0.2; adj-*P* >0.99; Figure 1F), ESP06B (χ^2^ _1_ = 0; adj-*P* >0.99; Figure 1H), Lw1047 (χ^2^ _1_ = 0; adj-*P* >0.99; Figure 1I), and PAO1 (χ^2^ _1_ = 0.2; adj-*P* >0.99; Figure 1J) showed similar virulence profile. Only strain AGO4092 showed a different virulence profile in the second trial, with a dip in survival happening after 72 hours in the first trial and after 96 hours in the second trial (χ^2^ _1_ = 25.8; adj-*P* < 0.001; Figure 1G). These results suggest that *C. elegans* while allowing for variance in virulence between strains, might be a more reliable model system between trials.

## 4. Discussion

Here, we show that passive exposure to an opportunistic pathogen in different animal hosts is a reliable model to investigate virulence variability and phenotypic. Specifically, we found that *P. aeruginosa* develops at least three virulence profiles when exposed to zebrafish larvae or developing *C. elegans*. Crucially, we also found that some strains of *P. aeruginosa* harbored different virulence profiles between the two models. The most striking example is strain A10, a clinical strain isolated from a wound in the early 20^th^ century, which incurred no significant mortality in zebrafish but ended up being one of the most virulent strains in the worm model. These results demonstrate that host-pathogen interactions depend not only on the host and the pathogen being considered but also on specific strains within a given pathogen. Our study thus highlights the need to consider genotypic and phenotypic diversity when investigating infectious disease and that using more than one model could enable to focus on different aspects of virulence in the bacteria.

To our knowledge, this is the first evidence that *P. aeruginosa* can incur mortality in zebrafish via passive exposure. While previous studies showed that *P. aeruginosa* PAO1 could induce mortality via microinjection ^30^, Soest et al. (2011) reported no mortality in larval zebrafish exposed to *P. aeruginosa* strain PAO1 for five hours post-fertilization. Here, we show that not only *P. aeruginosa* strain PAO1 can induce significant mortality in 5 dpf larvae. Here, we are confident that mortality in our study is predominantly attributable to the effect of the bacteria rather than other environmental variables such as oxygen depletion or pH changes due to bacterial growth ^42^ since our research clearly shows that bacterial growth does not always correlate with animal mortality. Similarly, using microscopy, we systematically screened for the presence of *Paramecium*, a common contamination of laboratory facilities feeding on dead animals ^43^, using microscopy. We found that *Paramecium* was only detectable visually 24 hours following a mortality incident, confirming that the parasite likely did not impact our study.

This study also confirms that *C. elegans* is a sound model system for investigating variance in virulence phenotype observed in *P. aeruginosa*. While the virulence of *P. aeruginosa* strain PAO1 was well documented in *C. elegans* ^34,35,44^, our study demonstrates that the nematode is sensitive enough to detect small differences in virulence between strains. Importantly, we used slow killing, caused by an active infection by live bacteria that accumulates in the lumen of the worm’s intestine ^33^. Thus, the time it took for all worms to die may be attributed to the virulence assay we used.

Our study also confirms that *P. aeruginosa* can infect a range of organisms, including zebrafish and *C. elegans* ^23^. Moreover, virulence shows a high level of heterogeneity between strains and, to a lesser extent, even within strain. While many reasons could explain these findings, differences in virulence traits and gene expression likely explain some of the changes between strains, and that gene expression explains, at least partly, changes within strains ^36,45,46^. In addition to changes in bacterial strains, changes in host immunity due to the brood effect or changes in gene expression could also affect the observed virulence phenotype ^47,48^, the latter potentially modulated by bacteria gene expression as well ^49^.

Another important consideration in our study is the role of temperature on virulence. According to the protocol for the SK assay detailed by Kirienko and colleagues, we incubated each strain at 37 °C for 24 hours and then moved our plates to 25 °C for another 24-hour incubation period to allow the bacteria to produce factors necessary for their full virulence ^33^. When assessing how *P. aeruginosa* adapts to temperature, researchers found that in PAO1 genes encoding enzymes needed for the biosynthesis of siderophores, pyochelin, and pyoverdine were regulated differently at 22 °C in comparison to 37 °C based on their transcriptome analysis ^50,51^. While they found that genes involved in pyoverdine biosynthesis and the pyoverdine operon regulator, ppyR, were upregulated at 22 °C, genes necessary for pyochelin biosynthesis were upregulated at 37 °C ^51^. This is important as the siderophore pyoverdine is a key virulence factor that provides *P. aeruginosa* with iron during infection, regulates the production of secreted toxins, as well as disrupts iron and mitochondrial homeostasis in the host ^52^. Thus, scientists should further investigate the role of temperature on the virulence of different *P. aeruginosa* strains in different model systems.

The ability to detect differences in larval zebrafish and *C. elegans* mortality between pathogenic strains provides an important model for studying virulence in opportunistic pathogens. Interestingly, we chose the five *P. aeruginosa* strains to represent a range of phenotypic diversity based on isolation site and date. While the size of our data set limits our ability to explore possible statistical associations between the different traits of the bacteria, we can conclude that there exists a range of discernable and repeatable virulence responses in the bacterium. The use of experimental models should enable the investigation of virulence evolution in a new light. Indeed, because the zebrafish adaptive immune system develops in zebrafish between two to four weeks post-fertilization ^53^, infection in larval zebrafish enables to focus on the virulence profile of the bacteria itself. For example, a high mortality rate quickly following exposure to bacteria could be explained by high levels of toxin expression or related to direct colonization of animal tissue ^54^. On the other hand, a lower mortality rate developing throughout exposure could result from virulence factors relating to bacterial growth or regulated via changes in gene expressions or quorum sensing ^55,56^. Further work is needed to identify the possible virulence mechanisms of each strain.

Taken together, our results suggest that passive exposure is a valuable tool to investigate the phenotypic diversity associated with host-parasite interactions in different model systems, offering the possibility to better understand pathogens’ virulence, including the emergence of novel pathogens from reservoir species.

## Acknowledgments

The authors thank Bard College for funding this project and Maureen O’Callaghan-Scholl for technical assistance with the zebrafish matings.

## Author Contributions

Conceptualization, D.A., A.E., H.B., and G.G.P.; methodology, D.A., A.E., M.T., M.O.S., H.B., and G.G.P.; writing—original draft preparation, D.A., and A.E.; writing—review and editing, H.B., and G.G.P..; supervision, H.B., and G.G.P.; funding acquisition, G.G.P.

## Conflicts of Interest

The authors declare no conflict of interest.

## Notes

### Competing Interest Statement

The authors have declared no competing interest.

